# Selective enrichment of specific bacterial taxa in downy mildew-affected spinach: Comparative analysis in laboratory and field conditions

**DOI:** 10.1101/2024.08.23.609345

**Authors:** Pim Goossens, Kim Baremans, Marrit Alderkamp, Jordi C. Boshoven, Guido van den Ackerveken, Roeland L. Berendsen

## Abstract

Plants host diverse microbial communities that can be influenced by their hosts to mitigate biotic stress. Previous research demonstrated that distinct laboratory cultures of *Hyaloperonospora arabidopsidis* (Hpa) on *Arabidopsis thaliana*, consistently harbor nearly identical bacteria. In this study, we analyzed the bacterial phyllosphere communities of laboratory-grown spinach plants infected by the downy mildew pathogen *Peronospora effusa* (Pe). Using 16S amplicon sequencing, we identified 14 Amplicon Sequence Variants (ASVs), with diverse taxonomies, that were enriched in at least 3 out of 5 investigated Pe cultures. This small set of 14 ASVs occupied on average 6.9% of the total bacterial communities in healthy spinach plants, and 43.1% in Pe-inoculated plants. A specific *Rhodococcus* and a *Paenarthrobacter* ASV were particularly prevalent and abundant. To validate these findings outside of the laboratory, we planted a susceptible variety of spinach in 4 agricultural fields and sampled leaves from Pe-infected plants in 2 fields where this pathogen naturally occurred. Comparative microbiome analysis of diseased and healthy plants revealed significant enrichment of 16 and 31 ASVs in these 2 fields, respectively. Among these, the *Paenarthrobacter* ASV was enriched in one field and the *Rhodococcus* ASV in the other field, suggesting that disease-associated microbiota that are abundantly detected in Pe laboratory cultures are also associated with Pe-infected field plants. Additionally, we observed an overlap of ASVs that were associated with both Pe and Hpa, indicating that similar bacteria are linked to downy mildew disease across different hosts.

## Introduction

Crop diseases cause global yield losses between 20% and 30% (Savary *et al*., 2019) and prevention strategies primarily rely on chemical pesticides and herbicides, or the use of resistant crop varieties obtained through breeding or genetic modification. However, crop protection chemicals pose risks to public health and the environment, and are increasingly subject to restrictive regulations (Barzman *et al*., 2015; Nishimoto, 2019). Moreover, pathogens can quickly evolve to break crop resistance (Stam *et al*., 2018). In contrast, the microbial communities associated with plants have long been recognized as a potential source of sustainable crop protecting agents (Bakker *et al*., 2020; Barratt *et al*., 2018; Weller, 2007).

Recent research has highlighted that both plants and pathogens can modulate microbial communities to their benefit. Plants are able to recruit beneficial microbes upon pathogen attack (Berendsen *et al*., 2018; Carrión *et al*., 2019; Yuan *et al*., 2018) and can actively shape their microbiome through exudation of primary and secondary metabolites (Pascale *et al*., 2020; Stringlis *et al*., 2018; Yu *et al*., 2021). Conversely, also pathogens have been shown to secrete effector proteins not only to suppress plant immunity, but also to antagonize their microbial antagonists (Snelders *et al*., 2020). A thorough understanding of such mechanisms driving microbiome assembly and the development of plant disease is crucial for developing innovative microbiome-based approaches to control plant disease.

Our previous research on *Arabidopsis thaliana* infected by the downy mildew *Hyaloperonospora arabidopsidis* (Hpa) revealed the consistent enrichment of specific disease-associated microbiota (Goossens *et al*., 2023). Hpa, an obligate biotroph, is maintained in culture on live host plants in several European research laboratories, where spores are successively washed from infected leaves and used to inoculate newly grown plants. Near-isogenic bacteria were found to accumulate in distinct Hpa cultures in labs in the Netherlands and Germany with >99.99% average nucleotide identity between the genomes of isolates obtained from these distinct Hpa cultures. These specific bacterial genomes were additionally identified in Hpa cultures of two distinct laboratories in the United Kingdom, based on metagenomic data. Interestingly, most of these bacteria negatively affect downy mildew proliferation (Goossens *et al*., 2023). Additionally, although the downy-mildew associated bacteria are particularly abundant on infected leaves, they can persist in soil, and form a soil-borne legacy (Bakker *et al*., 2018) that protects a next generation of plants growing on the soil (Berendsen *et al*., 2018; Goossens *et al*., 2023). Moreover, infection of Arabidopsis plants by gnotobiotic spores of Hpa resulted in specific promotion of downy-mildew associated bacteria from a diverse natural soil microbiome (Goossens *et al*., 2023).These findings suggest that plants, upon downy-mildew infection, can selectively promote specific bacteria that help protect against disease. However, it is unclear whether these downy mildew-associated microbiomes are unique to laboratory conditions or whether they also occur on field-grown downy-mildew-diseased plants.

While *Arabidopsis* and Hpa are well-studied model systems, spinach (*Spinacia oleracea)* and its cognate downy mildew, *Peronospora effusa* (Pe; previously *Peronospora farinosa f. sp. spinaciae*), represent an economically-relevant pathosystem. Downy mildew is the most important disease of spinach (Correll *et al*., 2011; Correll *et al*., 2007; Correll *et al*., 1994) with field-grown spinach plants having a high risk of infection by Pe often leading to harvest loss especially in organic production. Downy mildew-infected plants display yellowing of leaves corresponding to sporulation sites, where asexual conidia (spores) are produced on the leaf surface, capable of being dispersed by wind and water, causing secondary infections in the field (Correll *et al*., 1994).

Numerous races of Pe have been described, defined by their (in)compatibility with a set of differential spinach cultivars. As with Arabidopsis, differential compatibility is caused by presence or absence of specific resistance genes in plant cultivars (Correll *et al*., 2011). Pe races are cultivated for experimentation in successive cycles of reproduction on susceptible plants. Different Pe races have often been cultivated for extended periods of time in continuous isolation from each other. In this study, we investigated whether different Pe-infected plants carry specific disease-associated microbiomes, both in laboratory-maintained *in planta* Pe cultures and in naturally Pe-infected field-grown spinach.

## Materials and methods

### Plant growth conditions prior to inoculation

For the experiments at UU, 770-ml pots were filled with approximately 230 g of potting soil and saturated with water. Four *Spinacia oleracea* seeds of either cultivar Caladonia or Viroflay (kindly provided by Peter Paul Damen, Rijk Zwaan) were sown per pot approximately 0.5 cm below the soil surface and immediately placed in a climate-controlled chamber (21 °C, 70% relative humidity, 16 h light/8 h dark, light intensity 100 μmol m^−2^ s^−1^) where seeds were then allowed to germinate and develop. After germination, excess seedlings were removed to retain a single growing plant per pot. For the experiments at Location 2, Viroflay seeds were sown in a perlite-compost mix in 216 ml pots, 30 seeds per pot, and immediately placed in a climate-controlled chamber (21 °C, 65% relative humidity, 16 h light/8 h dark. For the experiments at Location 3, Viroflay seeds were sown in potting soil in 770 ml pots, 4 seeds per pot, and immediately placed in a climate-controlled chamber (23 °C, 16 h light/8 h dark, with extra lighting for 7h/day with assimilation lamps).

### Pe infection assays

Pe spore suspensions were prepared by vigorous shaking and subsequent vortexing (5 seconds) of infected shoot material (*i*.*e*. after removing plant roots by cutting the hypocotyl) in 50 ml tubes and diluted with water to obtain densities of 75 spores/μl.

For experiments performed at UU, 22-day-old spinach plants were spray-inoculated using paintbrush guns and pressurized air with spore suspensions in autoclaved tap water containing either Pe races Pe10, Pe11, or Pe14 (75 spores/μl). Wind inoculations of these races were performed by placing a pot with healthy plants below a Pe*-*inoculated and heavily*-*sporulating leaf that was held at a 90 degree angle. Spores were subsequently dispersed from the sporulating plants by a brief pulse of compressed air from above. Between 5 to 10 sporulating leaves were used for the wind-inoculation of 20 plants (10 Viroflay and 10 Caladonia). Control plants were left uninoculated. Pots were randomly placed in a climate chamber (16 °C, 10 h light/14 h dark, light intensity 100 μmol m−2 s−1), and were covered with moisturized transparent lids to increase humidity and covered by aluminium foil to keep the plants in the dark. This foil was removed after 24 hrs and trays were opened slightly to allow for ventilation. Samples were collected 12 days post-inoculation. For the experiment performed at Location 2, 9-day-old Viroflay seedlings were spray-inoculated with an excess of Pe14 spores in water or left uninoculated as controls. After spraying, plants were covered and kept in the dark at 100% relative humidity for 24hrs and placed into a climate-controlled chamber (16 h light/8 h dark, 16 °C / 14 °C. Samples were collected 10 days post-inoculation. At Location 3, 18-day-old spinach plants were spray-inoculated with Pe14 or left uninoculated and placed into a climate-controlled chamber (14 °C/12 °C, 14 h light/ 10 h dark, 100% relative humidity). Samples were collected 11 days post-inoculation.

### Spinach sample collection and genomic DNA extractions

For laboratory-grown spinach, 1 leaf per plant was collected, carefully avoiding the sampling of soil, in a 50-ml tube containing 3 glass beads, snap-frozen in liquid nitrogen, and stored at -80 °C until further processing. In the field, single leaves were collected, transported on ice and frozen at -80 °C upon arrival approximately 2 hours after harvest. Total genomic DNA (gDNA) was extracted from spinach leaves and resident microbiomes using the PowerLyzer PowerSoil DNA isolation kit (Qiagen) modified for leaf material, as described by Goossens *et al*. (2023). The quality and quantity of isolated gDNA was subsequently assessed using Nanodrop (Thermo Scientific).

### Pe quantification by qRT-PCR

To quantify Pe, two-step qRT–PCR reactions were performed in optical 96-well plates with a ViiA 7 real time PCR system (Applied Biosystems), using the Power SYBR® Green PCR Master Mix (Applied Biosystems) with 0.8 μmole L^-1^ primers that targeted Pe (this study) and spinach *ACTIN* (Duressa *et al*., 2012) (Supplementary Table S6). A standard thermal profile was used as described by Goossens *et al*. (2023). Pe abundance was calculated as 2^−(*CtPeACTIN*−*CtSpinachACTIN)*^.

### 16S rDNA amplicon library preparation for microbiome analysis

Library preparations for Illumina 16S rDNA amplicon sequencing with MiSeq V3 chemistry (2×300 base pairs paired-end sequencing) were performed on aforementioned gDNA samples based on standard Illumina protocols (illumina.com). Libraries were prepared using materials and methods as described by Goossens *et al*. (2023) with the use of primers (Supplementary Table S6) with heterogeneity spacers for the 16S amplicon PCR reaction (de Muinck *et al*., 2017). Sequencing was performed at the Utrecht Sequencing Facility (Useq; Utrecht, the Netherlands).

### 16S rDNA amplicon sequencing analyses

Preprocessing of sequencing data was performed as described by Goossens *et al*. (2023). Specific for the present study, ASVs were removed those that contributed to the lowest 1% (laboratory experiments), 10% (Warmenhuizen field samples), or 5% (Andijk field samples) of total cumulative abundances, based on the shape of the total cumulative abundance curve. These ASVs constitute the tail of the cumulative abundance curve that approached the plateau of that curve. Lastly, ASVs were removed that occurred in fewer than 6 samples.

All calculations of beta diversity, plots, and differential abundance tests were performed or made in R (version 4.2.0), with the exception of the Venn-diagram, for which the VIB webtool was used (http://bioinformatics.psb.ugent.be/webtools/Venn/). All PCoA ordinations and PERMANOVA tests were performed on Bray-Curtis dissimilarity matrices calculated for relative abundance data, using either vegan (version 2.6.4) or vegan functionalities within the phyloseq package (version 1.42.0). The pairwiseAdonis package (version 0.4) was employed for PERMANOVA tests involving multiple comparisons. Differential abundance testing was performed using ANCOM-BC (Lin *et al*., 2020) (script as specified at https://github.com/FrederickHuangLin/ANCOM-BC-Code-Archive/blob/master/scripts/ancom_bc.R). All graphs were made using ggplot2 (version 3.4.0) or ggpubr (version 0.5.0). Data wrangling was done with packages from the Tidyverse suite.

## Results

### Characterization of bacterial communities of Peronospora effusa-treated spinach leaves

To investigate Pe-associated microbiomes, we analyzed the bacterial communities of spinach leaves inoculated with *Peronospora effusa* (Pe) race Pe10, Pe11, or Pe14. These Pe races are routinely maintained in culture for research purposes by washing spores from infected leaves and subsequently spray inoculating the spores onto new susceptible plants. In this study, we used 2 spinach cultivars; Viroflay, which is susceptible, and Caladonia, which is resistant to all three Pe races. Plants were left uninoculated (controls) or were inoculated with Pe using two distinct methods to mimic the primary modes of Pe dispersal in the field. Leaves of Viroflay spinach plants infected by Pe with visible sites of spore production were washed in sterilized water to create spore suspensions. These suspensions were then used to spray-inoculate healthy plants (referred to as water inoculation). Alternatively, spores were transferred from a sporulating Viroflay leaf to adjacent healthy plants using a stream of compressed air (referred to as wind inoculation).

Eight days after inoculation, the first sporulation of Pe was observed on leaves of susceptible Viroflay plants only, regardless of whether plants were inoculated by wind or water. Pe abundance was quantified using real-time quantitative PCR (RT-qPCR; Fig. 1A, B) from genomic DNA extracted from leaves at twelve-days-post inoculation. All three Pe races successfully infected and colonized the susceptible Viroflay plants (Fig. 1A), but not the resistant Caladonia plants (Fig. 1B), regardless of the inoculation method. The inoculated samples and uninoculated controls were then processed for bacterial community analysis by next generation sequencing of 16S rDNA amplicons.

**Figure 1.**
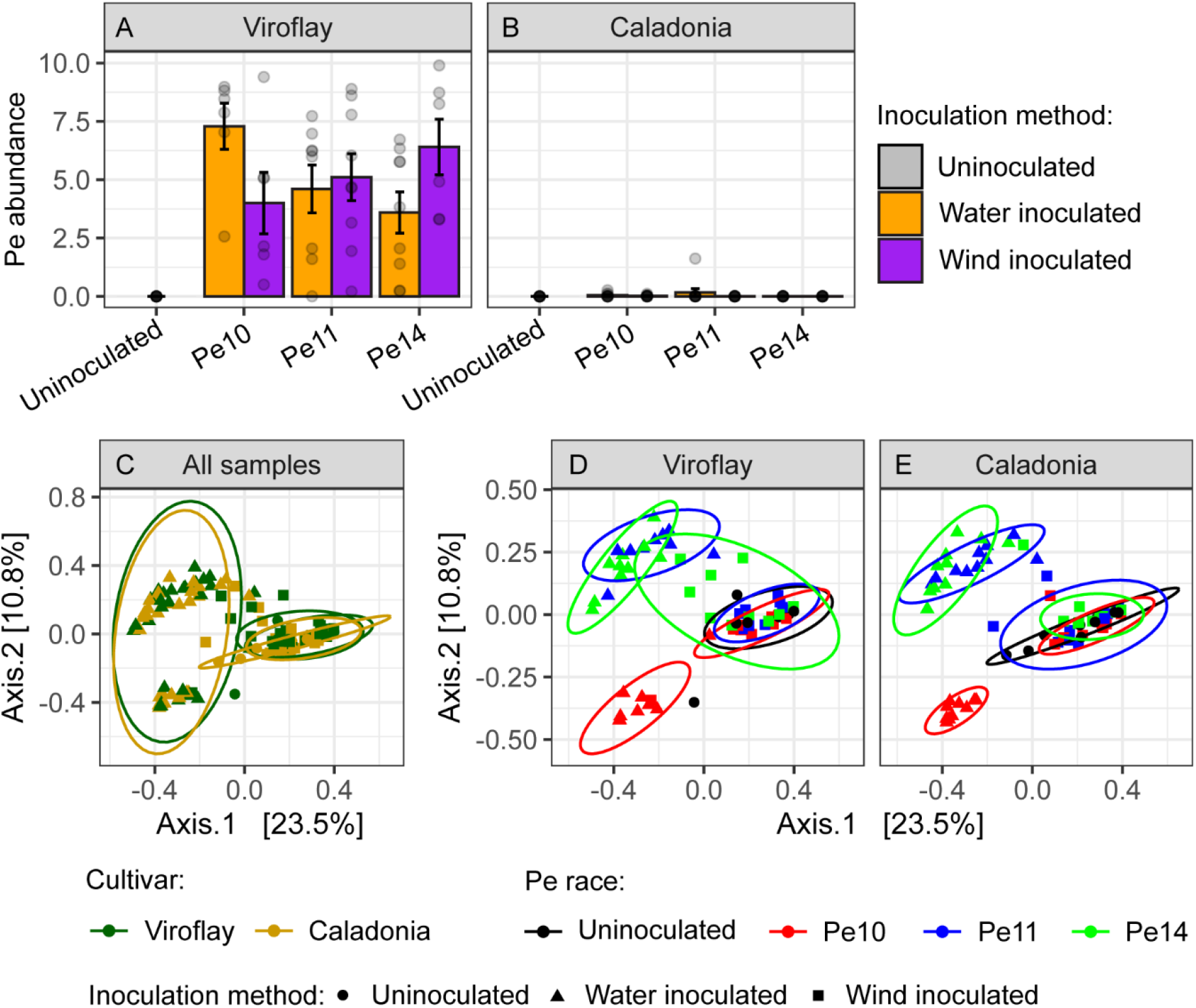
Pe infection and bacterial community shifts in spinach phyllospheres upon water- or wind-inoculation of susceptible (Viroflay) and resistant (Caladonia) cultivars. (**A, B**) Bar graph showing Pe abundance based on qPCR quantification of Pe genomic *ACTIN*, normalized by spinach genomic *ACTIN*, for (**A**) Viroflay and (**B**) Caladonia plants that were uninoculated or 12 days post-inoculation with Pe10, Pe11, or Pe14 by water- (orange bars) and wind inoculation (purple bars). Error bars show standard error of the mean (*N* = 10). The vertical axis was limited to 10.0: higher individual values are thus not visible in the figure. (**C** - **E**) PCoA ordination plots based on Bray-Curtis dissimilarities of uninoculated (circles), Pe water-inoculated (triangles), and Pe wind-inoculated (squares) plants. Samples are colored based on (**C**) cultivar or (**D, E**) treatment as indicated in the legends next to the ordination plots. (**D**) Viroflay and (**E**) Caladonia samples were subsetted from the same ordination space (**C**) to visualize the similarity in treatment-dependent shifts between susceptible and resistant host plants. Ellipses represent multivariate t-distributions with a 95% confidence level.

A total of 18,555,854 bacterial reads were generated, which, after quality filtering, were reduced to 14,841,570 reads comprising 921 amplicon sequence variants (ASVs) in 140 samples (*N* = 10 per treatment group; uninoculated Viroflay and Caladonia, Pe10, Pe11, and Pe14 water- or wind-inoculated Viroflay and Caladonia plants).

Spinach cultivar had a minor but significant effect on community structure in uninoculated leaf samples (*R*^2^=0.08, *P* = 0.0247), whereas these cultivar-dependent differences were mostly no longer significant after water or wind inoculation with Pe (Supplemental Table S1). This indicates that the significant minor shift in community structure for uninoculated samples is negated after Pe inoculation.

Upon water inoculation with Pe isolates Pe10, Pe11, and Pe14, substantial and significant shifts in community structure were observed compared to uninoculated samples for both susceptible (Viroflay) and resistant (Caladonia) cultivars (Fig. 1C - E, Supplemental Table S2). The phyllosphere community that developed on plants following water inoculation was distinct for each of the Pe races, yet similar on resistant and susceptible plants (Fig. 1D, E; Supplemental Table S1, S2, S3). This suggests that the majority of the community changes upon Pe water inoculation result from co-inoculation of bacteria in Pe spore suspensions, rather than the direct effect of Pe infection.

In contrast, Pe wind inoculation did not result in substantial community shifts (Fig. 1C - E; Supplemental Table S2). Only wind inoculation of Pe14 significantly affected community structure. Pe14 wind-inoculated Viroflay samples differed significantly from uninoculated samples, but also from Pe10 and Pe11 wind-inoculated Viroflay samples. These differences were not observed for the similarly-treated resistant Caladonia cultivar (Supplemental Table S2). This suggests that these smaller changes following wind inoculation are dependent on successful Pe infection and that few bacteria are transported along with the Pe spores during wind inoculation.

### Co-dispersal of bacteria alongside Pe spores occurs more readily in water than in wind

To identify bacterial taxa associated with Pe, we performed differential abundance testing for Viroflay samples using ANCOM-BC (Lin *et al*., 2020). For each of the three Pe isolates, more ASVs were significantly enriched in the phyllosphere of infected Viroflay plants following water inoculation (between 18-32 ASVs; Fig. 2A-C) than following wind-inoculation (between 4-5 ASVs Fig. 2A-C).

**Figure 2.**
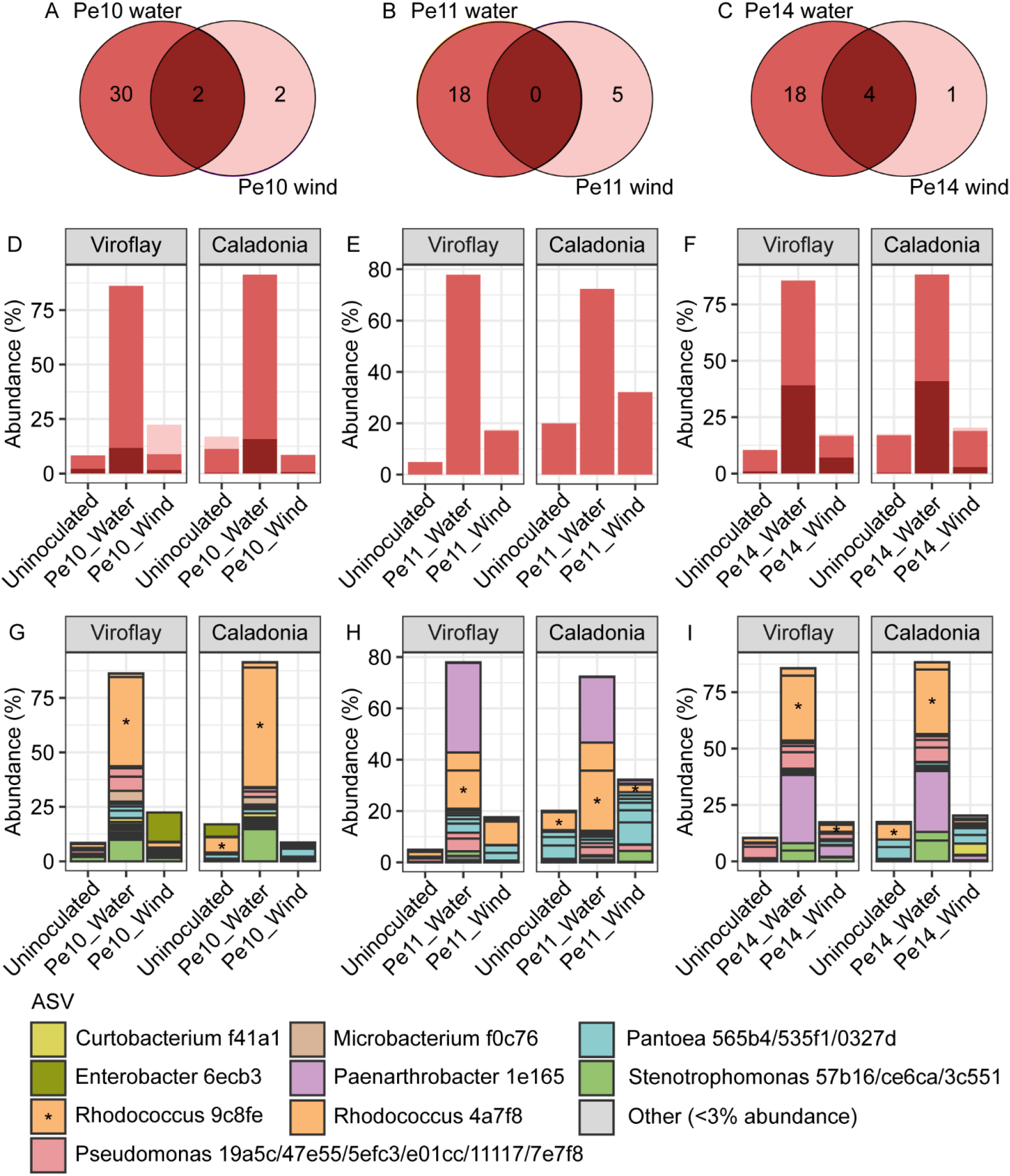
Significantly enriched ASVs in Pe water-inoculated Viroflay leaves are likely co-inoculated with Pe spores, whereas co-dispersal is more limited via wind-inoculation. (**A, B, C**) Venn diagrams of ASVs that were significantly enriched in Viroflay leaves inoculated water and wind with (**A**) Pe10, (**B**) Pe11, or (**C**) Pe14. (**D, E, F**) Bar graphs of the average cumulative abundances of ASVs that were significantly enriched in Viroflay leaves that were inoculated via water (medium red), wind (light red), or via both water and wind (dark red) with (**D**) Pe10, (**E**) Pe11, or (**F**) Pe14. (**G, H, I**) Bar graphs of the average abundances of individual ASVs that were significantly enriched in Viroflay leaves that were inoculated via water or wind with (**G**) Pe10, (**H**) Pe11, or (**I**) Pe14. (**D – I**) The abundances of these ASVs were visualized in both Viroflay (susceptible host; left panels) and Caladonia (resistant host; right panels) to illustrate that these bacteria are likely co-inoculated with Pe spores. (**G – I**) Colors represent taxonomy of individual ASVs as indicated below the Figure. Only ASVs that had an average abundance of at least 3% in any sample group were assigned colors. These ASVs are indicated in the color legend by 5-character codes. Significant enrichment is based on ANCOM-BC; Bonferroni-corrected *P* < 0.05 or structural zero ASV with presence in at least 5 more infected plant samples than in uninfected samples.

The ASVs that were enriched after water inoculation dominated the phyllosphere bacterial communities and together comprised more than 70% of the total communities in these samples, whereas the cumulative abundance of these same ASVs was below 12% in uninoculated plants (Fig. 2D-F). This pattern was observed for both susceptible Viroflay plants and resistant Caladonia plants, further indicating that these bacteria are likely co-inoculated with Pe spores suspensions obtained from infected leaves. As the Pe water inoculation method used in this study is identical to the inoculation method used in routine *in planta* maintenance of Pe in the laboratory, these bacteria are likely to persist in Pe laboratory cultures, due to co-dispersal with each passage of the pathogen to new host plants.

Numbers of enriched ASVs following Pe wind inoculation were more limited (Fig. 2A - C) and the average abundances of the enriched ASVs were generally lower than following water inoculation (Fig. 2D - F; Extended data 2). Some overlap was observed between the ASVs that were enriched after water and wind inoculation of Pe10 (Fig. 2A) and Pe14 (Fig. 2C). The enrichment of Pe-associated ASVs after wind-inoculation was most evident for Pe14 (Fig. 2F). This is in line with the abovementioned PERMANOVA tests in which only Pe14 wind-inoculated Viroflay samples differed significantly from uninoculated Viroflay samples (Supplemental Table S2).

Upon inspection of the taxonomy of ASVs enriched following inoculation, two genera appear to strongly dominate the phyllospheres of particularly Pe water-inoculated samples. Water-inoculated samples contained high abundances of *Rhodococcus* ASVs regardless of plant genotype or Pe race (Fig. 2G-I) and exhibited particularly high loads of *Rhodococcus* ASV 9c8fe (Fig 2G-I; Extended Data 2). This ASV occupied between 15% and 55% of Pe water-inoculated leaves on average, whereas its abundance was 1.9% in uninoculated Viroflay plants and 7% in uninoculated Caladonia plants. Additionally, in Pe11 and Pe14 water-inoculated phyllospheres a single *Paenarthrobacter* ASV (1e165) occurred in high abundance (Fig. 2H, I; Extended Data 2). In these water-inoculated treatments, *Paenarthrobacter* ASV 1e165 occupied on average between 25% and 35% of the total bacterial communities. Strikingly, this ASV was also one of the 5 ASVs that was significantly enriched in Pe14 wind-inoculated Viroflay samples (Extended Data1, Extended Data 2). There it accounted for on average approximately 5% of the total communities, whereas its abundance was 0.6% on average in uninoculated Viroflay samples. These findings suggests that *Paenarthrobacter* ASV 1e165 is promoted in the phyllosphere due to Pe infection.

*Rhodococcus* ASV 9c8fe and *Paenarthrobacter* ASV 1e165 are thus not only enriched and highly abundant in Pe infected leaves, they are also shared by cultures of distinct and separately-maintained Pe races. Moreover, these bacteria will likely persist in these cultures as they are co-dispersed with Pe during routine water-mediated passaging of the pathogen to new host plants.

### Identical bacterial ASVs are enriched in Pe cultures at separate locations

The dominance of bacteria that produce the exact same ASVs in cultures of distinct Pe races suggests that there are very specific selective pressures in Pe-infected phyllospheres. However, for the experiment above, the Pe10, Pe11 and Pe14 cultures had been brought to a single location (Utrecht University). To determine if the enrichment of these specific ASV also associated with Pe-infected plants at other locations, we sampled water-inoculated and uninoculated Viroflay control plants at the research facilities of two spinach-breeding companies. Cultures of Pe14 have been routinely maintained at these two companies since the denomination of this isolate in 2012. Since then, these have been cultured separately from each other as well as from the culture used in the experiments above (Pe14-1) which was kindly provided by a third breeding company.

In total, we sampled the phyllospheres of 40 plants for 16S amplicon sequencing (*N* = 10, 2 locations, uninoculated and water-inoculated), generating 4,712,621 bacterial reads that were reduced by filtering to 3,197,840 reads comprising 848 ASVs.

At both additional locations, water inoculation of Pe resulted in clear shifts in microbial communities leading to a phyllosphere bacterial community that was significantly different from control plants (Fig. S1A, B). We then again used differential abundance testing to identify bacterial ASVs that were significantly more abundant on Pe14-inoculated plants than uninoculated control plants at each location (Extended data 1). By cross-referencing the enriched ASVs from the Pe cultures at locations 2 and 3 (Pe14-2 and Pe14-3, respectively) with the Pe10, Pe11 and Pe14-1 cultures at Utrecht University, we found that there is indeed overlap at the ASV level between these cultures (Table 1; Fig. 3A). We found that 14 ASVs were significantly enriched in more than 3 out of 5 Pe cultures, 4 of which were enriched in 4 out of 5 Pe cultures, and 1 ASV was enriched in all Pe cultures that were investigated. The 14 ASVs that were enriched in at least three out of five Pe cultures occupied on average 6.9% of the total community in uninoculated Viroflay samples and 43.1% in Pe inoculated Viroflay samples, showing that this small set of ASVs commonly dominated the phyllospheres of Pe-inoculated plants (Extended Data 3; Fig. 3B). The consistent enrichment of specific ASVs in distinct Pe cultures across multiple locations indicates that specific bacteria are commonly favored in Pe cultures, and that these ASVs are not specific to the culture conditions, methods, and materials in our lab.

**Table 1.** ASVs consistently associated with Pe in 5 examined laboratory Pe cultures. ASVs that were significantly enriched in at least 3 separate Pe cultures are shown. Microbiomes of susceptible Viroflay plants that were water-inoculated with Pe10, Pe11, or Pe14 were compared to their uninoculated controls. Pe14 cultures were examined at 3 distinct locations (Pe14-1, Pe14-2, and Pe14-3, respectively). Significant enrichment is based on ANCOM-BC; Bonferroni-corrected *P* < 0.05 or structural zero ASV with presence in at least 5 more infected plant samples than in uninfected plant samples.

| ASV | Enriched in number of cultures | Enriched following water-inoculation with |
| --- | --- | --- |
| *Rhodococcus* ASV 9c8fe | 5/5 | Pe10; Pe11; Pe14-1; Pe14-2; Pe14-3 |
| *Rhodococcus* ASV 4a7f8 | 4/5 | Pe10; Pe11; Pe14-1; Pe14-2 |
| *Curtobacterium* ASV f41a1 | 4/5 | Pe10; Pe11; Pe14-1; Pe14-3 |
| *Stenotrophomonas* ASV ce6ca | 4/5 | Pe11; Pe14-1; Pe14-2; Pe14-3 |
| *Paenarthrobacter* ASV 1e165 |  |  |
| *Pantoea* ASV 0327d | 3/5 | Pe10; Pe11; Pe14-3 |
| *Pantoea* ASV 481d0 |  |  |
| *Pseudomonas* ASV 47e55 | 3/5 | Pe10; Pe14-1; Pe14-3 |
| *Rhodococcus* ASV ca9c4 |  |  |
| *Microbacterium* ASV f0c76 |  |  |
| *Brevundimonas* ASV 2cd30 |  |  |
| *Sphingobium* ASV ed6be | 3/5 | Pe10; Pe14-2; Pe14-3 |
| *Pantoea* ASV 535f1 | 3/5 | Pe11; Pe14-1; Pe14-3 |
| *Pantoea* ASV 565b4 |  |  |

**Figure 3.**
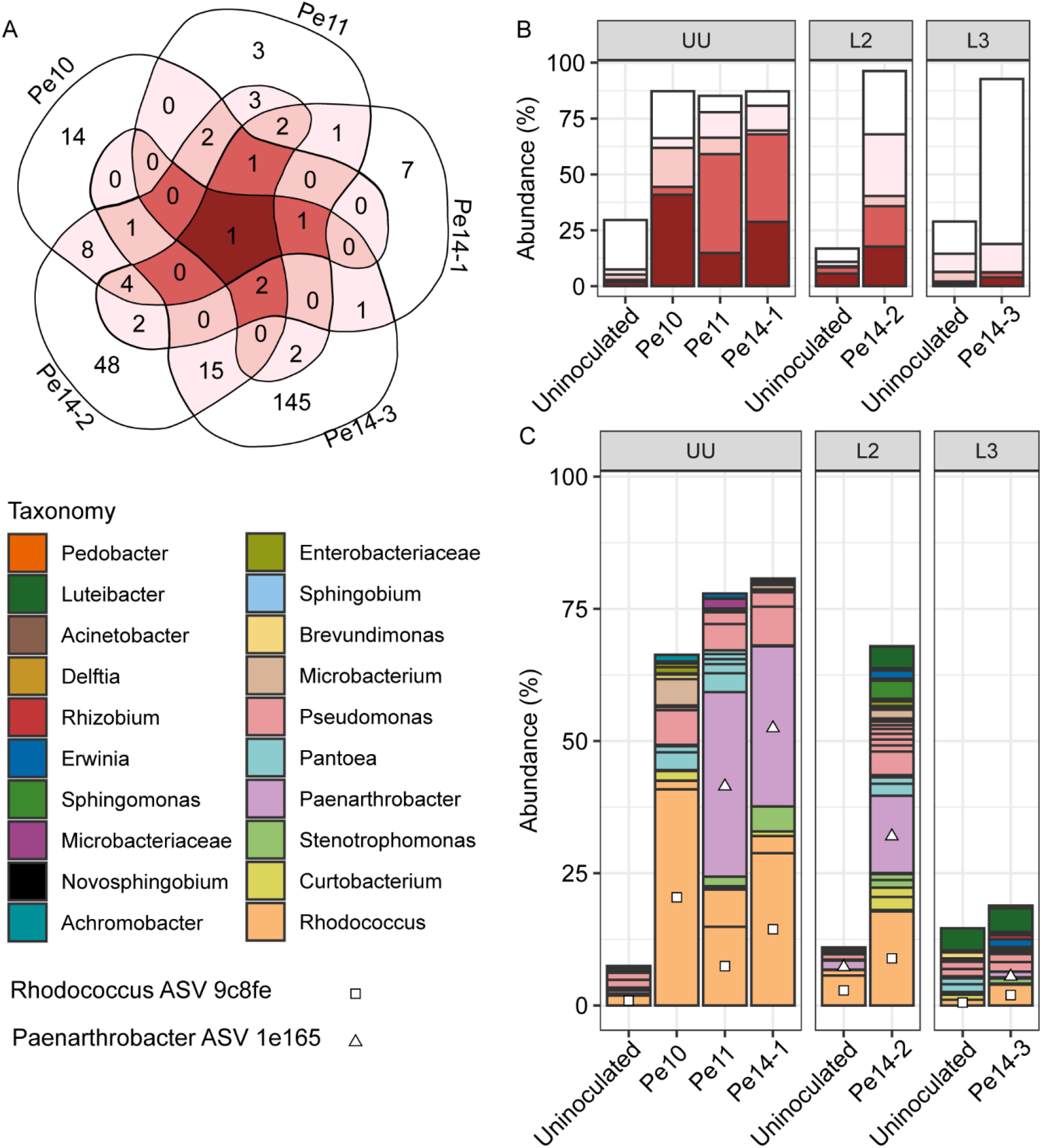
Phyllospheres of *Pe* cultures are generally dominated by a few bacterial taxa. (**A**) Venn diagram with all ASVs that were significantly enriched in at least one investigated *Pe* culture. (**B**) Bar graph showing the cumulative abundances of ASVs that were enriched in 1, 2, 3, 4, or 5 out of 5 *Pe* cultures, colored to match the Venn diagram. (**C**) Bar graph showing the abundances of individual ASVs that were significantly enriched in at least 2 out of 5 investigated *Pe* cultures. Colors indicate taxonomy as displayed on the left side of the graph. Squares indicate the *Rhodococcus* ASV 9c8fe and triangles indicate *Paenarthrobacter* ASV 1e165. The order of the individual ASVs in (**C**) approximately matches the order of the agglomerated ASVs in (**B**), although there are some discrepancies as the ASVs in (**C**) were ordered based on Taxonomy.

Plants water-inoculated with Pe14 at location 2 (Pe14-2) developed a bacterial community enriched by many ASVs shared with the Pe10, Pe11 and Pe14-1 cultures and to similar abundance levels as the Pe water inoculated plants at Utrecht University (Pe14-1; Fig. 3B, C). In the Pe14 culture at location 2 again *Rhodococcus* ASV 9c8fe and *Paenarthrobacter* ASV 1e165 were prevalent and abundant (Fig. 3C).

The phyllosphere bacterial community profile of Pe-inoculated plants at location 3 (Pe14-3) was more distinct from the other 4 cultures (Fig. 3B, C). The majority of ASVs that were significantly more abundant in water-inoculated plants compared to uninoculated plants at location 3 were not enriched in the other 4 cultures. Nonetheless, *Rhodococcus* ASV 9c8fe and *Paenarthrobacter* ASV 1e165 were also significantly enriched in Pe14-3-inoculated plants compared to uninoculated controls (Fig. 3C; Table 1; Extended data 3).

The investigation of the bacterial communities in separate Pe laboratory cultures thus reveals that these systems are enriched for specific bacterial taxa, defined at the ASV level. Specific *Rhodococcus* and *Paenarthrobacter* spp. commonly dominate the phyllosphere communities of Pe-infected spinach leaves, and bacterial ASVs corresponding to *Curtobacterium, Stenotrophomonas, Pantoea, Sphingobium, Pseudomonas, Microbacterium*, and *Brevundimonas* spp. are also enriched in Pe-infected spinach with varying degrees of consistency (Table 1).

### Natural downy-mildew infection in field-grown spinach

Although data from *Pe* laboratory cultures show consistent enrichment of specific ASVs, we cannot rule out that this may result from laboratory practices rather than from downy mildew infection itself. To test whether similar bacteria associate with downy mildew under natural conditions, we planted the susceptible spinach cultivar Viroflay in four trial fields in the Netherlands, near the villages of Warmenhuizen, Andijk, Schaarsbergen, and De Lier, and monitored disease symptoms. While *Pe* was not observed in Schaarsbergen, it did appear in Warmenhuizen, Andijk, and De Lier. In De Lier, infection levels were already high at the time of detection, preventing the sampling of symptomless plants. However, in Warmenhuizen and Andijk, we sampled leaves from both uninfected and *Pe*-infected plants showing symptoms like leaf yellowing and grey sporulation.

We quantified *Pe* abundance in these samples using qPCR on genomic DNA from Viroflay leaves, confirming that most symptomatic plants were colonized by *Pe*, while most asymptomatic plants were not. Pe, however, was detected in two plants from the Warmenhuizen field that were previously considered symptomless and these plants were reclassified as infected (Supplemental Fig. S2A). Similarly, four yellowing plants from the Andijk field were reclassified as uninfected, as Pe was not detected in these samples (Supplemental Fig. S2C). Pe abundance varied strongly between the two locations, with lower infection levels in Warmenhuizen compared to Andijk (Supplemental Fig. S2B, D).

### Pe-associated bacterial ASVs are shared between lab and field

To identify bacteria associated with Pe in the field, genomic DNA samples were processed for 16S rDNA amplicon sequencing. Sequencing generated 1,953,783 bacterial reads that were reduced to 1,658,419 reads comprising 1,838 ASVs after filtering in 24 Warmenhuizen samples, and 1,983,583 bacterial reads that were reduced to 1,799,117 reads comprising 836 ASVs in 28 Andijk samples.

At the community level, the phyllosphere microbiome showed no significant difference between uninfected and infected samples from Warmenhuizen (*R*^2^ = 0.04, *P* = 0.4; Fig. 4A). However, the samples with higher Pe infection levels from the Andijk field did reveal clear community differentiation (*R*^2^ = 0.35, *P<*0.001 in PERMANOVA; Fig. 4B).

**Figure 4.**
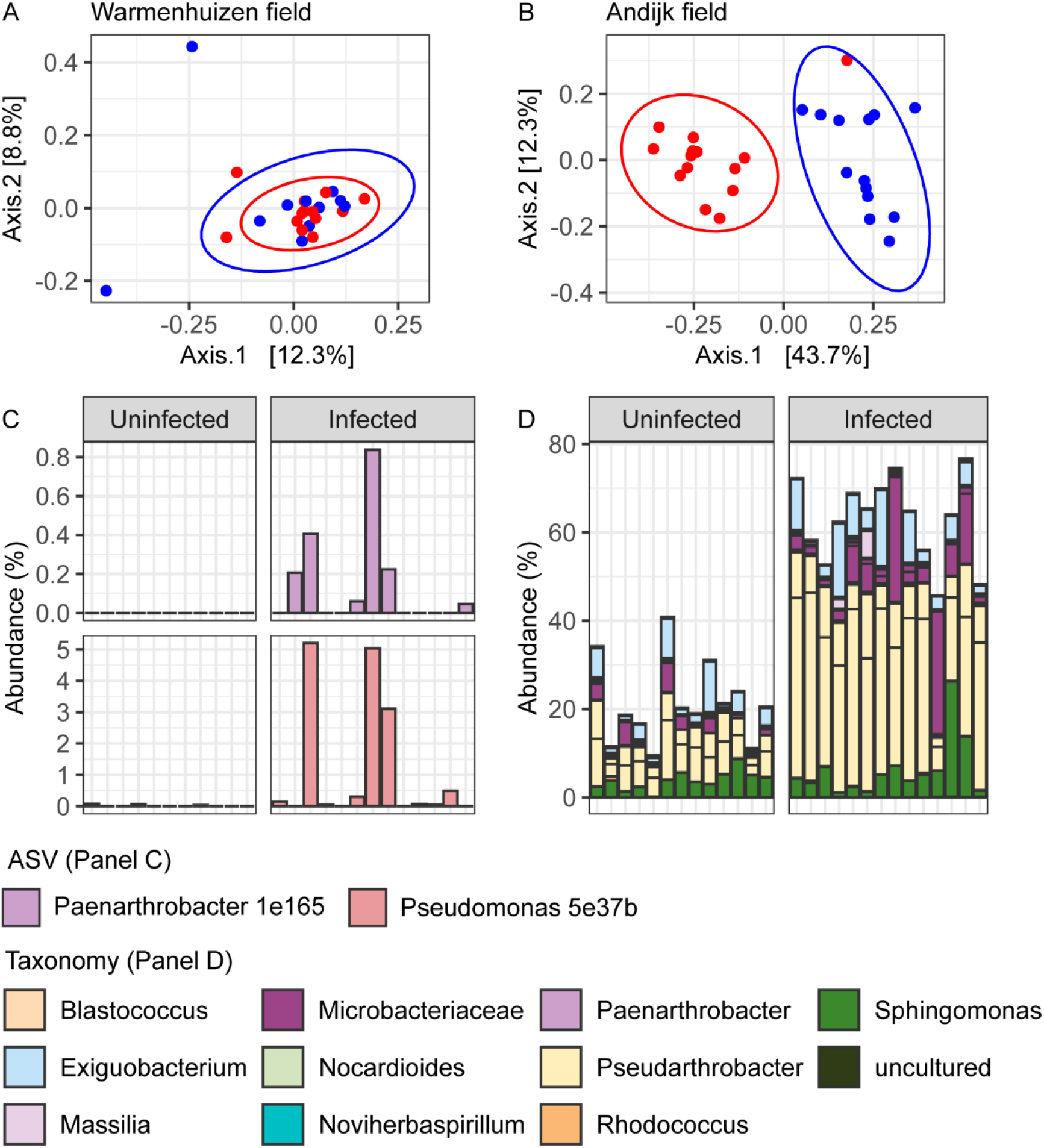
Pe-associated bacteria in field grown spinach. (**A, B**) PCoA ordination plots based on Bray-Curtis dissimilarities of uninfected (blue symbols) and *Pe*-infected (red symbols) Viroflay plants in fields at (**A**) Warmenhuizen and (**B**) Andijk. Text below the graphs show PERMANOVA results, per location, for the comparison between *Pe*-infected and uninfected plants based on 9999 permutations. (**C**) Bar graph showing the abundances of *Paenarthrobacter* 1e165 and *Pseudomonas* 5e37b per sample in uninfected and *Pe*-infected leaves in field-grown Viroflay plants at Warmenhuizen. (**D**) Stacked bar graph showing the abundances of individual ASVs that were significantly enriched in Pe-infected leaves per sample from the field at Andijk. ASV are colored based on taxonomy at the genus level as indicated on the right side of the graph. Significant enrichment for field data is based on ANCOM-BC; Benjamini-Hochberg-corrected *P* < 0.05 or structural zero ASV with presence in at least 6 more infected plant samples than in uninfected samples.

Although microbiome differentiation in Warmenhuizen was not significant at the community level, we detected 16 ASVs that were significantly enriched in Pe-infected leaves (Supplementary Table S4; Supplemental Fig. S3; Extended Data 1). These ASVs accounted for 2.52% relative abundance in Pe-infected samples, whereas they account for 0.55% of the communities in uninfected samples (Supplementary Table S4). Among these ASVs, we notably observe *Paenarthrobacter* ASV 1e165 and *Pseudomonas* ASV 5e37b, that were also enriched in lab cultures (Fig. 4C). *Paenarthrobacter* ASV 1e165 was observed in 6 out of 13 Pe-infected samples, but in none of the uninfected samples. Similarly *Pseudomonas* 5e37b was hardly detected in the uninfected samples, but abundant in 5 of the 13 infected plant samples (Fig. 4C).

In Andijk, differential abundance testing identified 31 ASVs significantly enriched in *Pe*-infected spinach leaves, comprising 63% of the total community in infected plants, compared to 21% in uninfected samples (Fig. 4D; Supplementary Table S5).(Fig. 4D; Supplementary Table S5). The high abundances of these enriched ASVs reflect the strong community-level differentiation between infected and uninfected samples in PCoA ordination (Fig. 4B). In Pe-infected samples, particularly high abundances were observed for *Exiguobacterium* 41452, *Pseudarthrobacter* c1d8f, *Pseudarthrobacter* d1316, *Sphingomonas* f359d, and *Microbacteriaceae* 8fc0f (Supplementary Table S5). Of these highly abundant ASVs, *Sphingomonas* f359d, and *Microbacteriaceae* 8fc0f were also enriched in some of the Pe lab cultures. Moreover, *Rhodococcus* 9c8fe, which was significantly enriched in five out of five investigated Pe laboratory cultures (Fig. 3), was also significantly enriched on leaves of infected plants in the Andijk field (Supplementary Table S5). Although its enrichment was not immediately evident from relative abundance (Supplemental Fig. S4; Supplementary Table S5), ANCOM-BC adjusted for sampling bias, revealing a significant increase in its log-transformed mean absolute abundance from 2.16 in uninfected samples to 3.81 in infected leaves at Andijk (Extended Data 4).

Together, data from lab and field experiments reveal a remarkably high co-occurrence of specific bacterial taxa, at the ASV level, with Pe. The data from Andijk also demonstrate that strong Pe infection can significantly alter phyllosphere microbiome composition in the field.

## Discussion

This study investigated the microbiomes of downy mildew-infected spinach leaves in both laboratory cultures of *Peronospora effusa* (Pe), and naturally-infected field-grown spinach. Similar to our observation for Hpa-associated microbiomes (Goossens *et al*., 2023), distinct and separately-maintained laboratory cultures of Pe share a small set of bacteria that are highly abundant on leaves of plants that are inoculated with these cultures, and absent or much less abundant in uninoculated control plants. In our study, 14 ASVs were each enriched in at least three out of five Pe cultures, and collectively accounted for nearly half of the total bacterial population. The consistent enrichment of these ASVs in both lab cultures and naturally infected field spinach suggests that specific microbes are promoted on Pe-infected plants due to particular selective cues that operate across different locations.

The enrichment of identical ASVs in distinct Pe cultures is highly unlikely to be coincidental. Prokaryotic diversity based on the 16S rDNA V4 region is estimated at 0.8 – 1.6 million operational taxonomic units (OTUs) (Louca *et al*., 2019) and alpha diversity estimates based on OTUs are largely equivalent to estimates based on ASVs (Glassman *et al*., 2018). Even if we assume that only 5000 ASVs can occur in the phyllospheres of spinach plants (very likely an underestimation), of which only 1000 ASVs occur in any random sample of 20 spinach plants, and of which only 50 ASVs (an overestimation) are significantly enriched in a set of 10 spinach plants versus a set of 10 other spinach plants, then the probability that 1 specific ASV is enriched in 5 such sets of spinach plants by random chance is 1^-10^. The probability that the enrichment of *Rhodococcus* ASV 9c8fe is coincidental in the cultures of Pe10, Pe11 and three separate cultures of Pe14, as well as in naturally Pe-infected field-grown spinach plants, is therefore negligible.

Although ASVs represent closely related bacteria and can distinguish different lineages within a species, a single ASV may still represent bacteria with different genomes and functions (Berry *et al*., 2017). Nonetheless, the Pe-associated bacteria represented by each ASV must at least share a larger set of genomic traits that provide them with the selective advantage that led to their enrichment. Previous work has shown that disease-associated bacteria represented by the same ASV, isolated from distinct Hpa cultures, are isogenic (Goossens *et al*., 2023), suggesting that the Pe-associated ASVs may also largely represent single bacterial genotypes.

Among the core members of the Pe laboratory microbiomes, that were significantly enriched in more than half of the Pe cultures (Table 1), we observed *Sphingobium* ASV ed6be and *Microbacterium* ASV f0c76. These ASVs were also previously associated with Hpa, the downy mildew of Arabidopsis (Goossens *et al*., 2023). *Sphingobium* ASV ed6be consistently appears in Hpa-associated microbiomes, while *Microbacterium* ASV f0c76, although enriched in only one Hpa culture, showed a decrease in abundance in the absence of Hpa, indicating a benefit from Hpa infection. The enrichment of these ASVs in the microbiomes of two distinct downy mildews is noteworthy.

Interestingly, a bacterium isolated from Hpa-infected leaves representing *Microbacterium* ASV f0c76 reduced Hpa proliferation when co-inoculated when co-inoculated with Hpa on Arabidopsis (Goossens *et al*., 2023). It is therefore tempting to speculate that *Microbacterium* ASV f0c76 but also other Pe-associated ASV identified in this study may similarly reduce downy mildew disease in spinach, akin to Hpa-associated microbiota that protect Arabidopsis. We hypothesize that plants may promote specific microbes in response to pathogen attack to defend against disease, a mechanism we are currently investigating.

In conclusion, our results show that the specific bacteria that are consistently enriched in laboratory Pe cultures also accumulate on Pe-infected spinach plants in the field. Our previous work with downy mildew-associated microbiota in *Arabidopsis* has shown that these microbes can protect plants from disease. Consequently, downy mildew maintenance cultures may harbor microbes with high potential for biological control, capable of colonizing plants most effectively when they are infected. Regardless, these microbes can be used to understand how disease-associated microbiomes are assembled and how they affect disease incidence and severity. Further investigations into the effects of these communities on Pe infection may reveal novel ways to combat downy mildew

## Acknowledgements

This project was supported by Topsector Horticulture & Starting Materials (project code 1605-106), with contributions by Rijk Zwaan, Bejo Zaden B.V., Pop Vriend Seeds, and DSM. We thank Johan Rijk for providing Pe isolates for starting cultures in our laboratory, and Annemiek Andel and Joyce Elberse for maintaining *in planta* cultures of Pe.

## Data availability statement

The data that support the findings of this study and isolates are available from the corresponding author upon reasonable request. Moreover, the raw amplicon sequence data generated in this study are available at https://www.ncbi.nlm.nih.gov/bioproject/PRJNA1106431 (samples from laboratory experiments) and at https://www.ncbi.nlm.nih.gov/bioproject/PRJNA1150921 (field samples).

## Author contributions

P.G., R.L.B., and G.A. designed experiments; P.G., K.B., M.A., and J.B. performed experiments; P.G. analyzed data; and P.G., R.L.B., and G.A. wrote the manuscript.

## Literature

Bakker, P. A., Berendsen, R. L., Van Pelt, J. A., Vismans, G., Yu, K., Li, E., Van Bentum, S., Poppeliers, S. W., Gil, J. J. S., & Zhang, H. (2020). The soil-borne identity and microbiome-assisted agriculture: looking back to the future. Molecular Plant, 13(10), 1394–1401.

Bakker, P. A., Pieterse, C. M., de Jonge, R., & Berendsen, R. L. (2018). The soil-borne legacy. Cell, 172(6), 1178–1180.

Barratt, B., Moran, V., Bigler, F., & Van Lenteren, J. (2018). The status of biological control and recommendations for improving uptake for the future. BioControl, 63(1), 155–167.

Barzman, M., Bàrberi, P., Birch, A. N. E., Boonekamp, P., Dachbrodt-Saaydeh, S., Graf, B., Hommel, B., Jensen, J. E., Kiss, J., & Kudsk, P. (2015). Eight principles of integrated pest management. Agronomy for sustainable development, 35(4), 1199–1215.

Berendsen, R. L., Vismans, G., Yu, K., Song, Y., de Jonge, R., Burgman, W. P., Burmølle, M., Herschend, J., Bakker, P. A., & Pieterse, C. M. (2018). Disease-induced assemblage of a plant-beneficial bacterial consortium. The ISME journal, 12(6), 1496–1507.

Berry, M. A., White, J. D., Davis, T. W., Jain, S., Johengen, T. H., Dick, G. J., Sarnelle, O., & Denef, V. J. (2017). Are oligotypes meaningful ecological and phylogenetic units? A case study of Microcystis in freshwater lakes. Frontiers in microbiology, 8, 365.

Carrión, V. J., Perez-Jaramillo, J., Cordovez, V., Tracanna, V., De Hollander, M., Ruiz-Buck, D., Mendes, L. W., Van Ijcken, W. F., Gomez-Exposito, R., & Elsayed, S. S. (2019). Pathogen-induced activation of disease-suppressive functions in the endophytic root microbiome. Science, 366(6465), 606–612.

Correll, J., Bluhm, B., Feng, C., Lamour, K., Du Toit, L., & Koike, S. (2011). Spinach: better management of downy mildew and white rust through genomics. European Journal of Plant Pathology, 129(2), 193–205.

Correll, J., Feng, C., Irish, B., Koike, S., Morelock, T., Bentley, T., & Tomlinson, A. (2007). Spinach downy mildew: overview of races and the development of molecular markers linked to major resistance genes. Advances in downy mildew research, 3, 135–142.

Correll, J. C., Morelock, T. E., Black, M. C., Koike, S. T., Brandenberger, L. P., & Dainello, F. J. (1994). Economically important diseases of spinach. Plant disease, 78(7), 653–660.

de Muinck, E. J., Trosvik, P., Gilfillan, G. D., Hov, J. R., & Sundaram, A. Y. (2017). A novel ultra high-throughput 16S rRNA gene amplicon sequencing library preparation method for the Illumina HiSeq platform. Microbiome, 5(1), 1–15.

Duressa, D., Rauscher, G., Koike, S. T., Mou, B., Hayes, R. J., Maruthachalam, K., Subbarao, K. V., & Klosterman, S. J. (2012). A real-time PCR assay for detection and quantification of Verticillium dahliae in spinach seed. Phytopathology, 102(4), 443–451.

Glassman, S. I., & Martiny, J. B. (2018). Broadscale ecological patterns are robust to use of exact sequence variants versus operational taxonomic units. MSphere, 3(4), 10.1128/msphere. 00148–00118.

Goossens, P., Spooren, J., Baremans, K. C., Andel, A., Lapin, D., Echobardo, N., Pieterse, C. M., Van den Ackerveken, G., & Berendsen, R. L. (2023). Obligate biotroph downy mildew consistently induces near-identical protective microbiomes in Arabidopsis thaliana. Nature Microbiology, 8(12), 2349–2364.

Lin, H., & Peddada, S. D. (2020). Analysis of compositions of microbiomes with bias correction. Nature communications, 11(1), 3514.

Louca, S., Mazel, F., Doebeli, M., & Parfrey, L. W. (2019). A census-based estimate of Earth’s bacterial and archaeal diversity. PLoS biology, 17(2), e3000106.

Nishimoto, R. (2019). Global trends in the crop protection industry. Journal of pesticide science, D19–101.

Pascale, A., Proietti, S., Pantelides, I. S., & Stringlis, I. A. (2020). Modulation of the root microbiome by plant molecules: the basis for targeted disease suppression and plant growth promotion. Frontiers in Plant Science, 10, 1741.

Savary, S., Willocquet, L., Pethybridge, S. J., Esker, P., McRoberts, N., & Nelson, A. (2019). The global burden of pathogens and pests on major food crops. Nature Ecology & Evolution, 3(3), 430–439.

Snelders, N. C., Rovenich, H., Petti, G. C., Rocafort, M., van den Berg, G. C., Vorholt, J. A., Mesters, J. R., Seidl, M. F., Nijland, R., & Thomma, B. P. (2020). Microbiome manipulation by a soil-borne fungal plant pathogen using effector proteins. Nature Plants, 6(11), 1365–1374.

Stam, R., & McDonald, B. A. (2018). When resistance gene pyramids are not durable—the role of pathogen diversity. Molecular plant pathology, 19(3), 521.

Stringlis, I. A., Yu, K., Feussner, K., De Jonge, R., Van Bentum, S., Van Verk, M. C., Berendsen, R. L., Bakker, P. A., Feussner, I., & Pieterse, C. M. (2018). MYB72-dependent coumarin exudation shapes root microbiome assembly to promote plant health. Proceedings of the National Academy of Sciences, 115(22), E5213–E5222.

Weller, D. M. (2007). Pseudomonas biocontrol agents of soilborne pathogens: looking back over 30 years. Phytopathology, 97(2), 250–256.

Yu, K., Stringlis, I. A., van Bentum, S., de Jonge, R., Snoek, B. L., Pieterse, C. M., Bakker, P. A., & Berendsen, R. L. (2021). Transcriptome Signatures in Pseudomonas simiae WCS417 Shed Light on Role of Root-Secreted Coumarins in Arabidopsis-Mutualist Communication. Microorganisms, 9(3), 575.

Yuan, J., Zhao, J., Wen, T., Zhao, M., Li, R., Goossens, P., Huang, Q., Bai, Y., Vivanco, J. M., & Kowalchuk, G. A. (2018). Root exudates drive the soil-borne legacy of aboveground pathogen infection. Microbiome, 6(1), 1–12.

