## Supplemental information for "Selective enrichment of specific bacterial taxa in downy mildew-affected spinach: Comparative analysis in laboratory and field conditions"

### Supporting information

Article title: **Consistent enrichment of specific bacterial taxa in spinach downy-mildew grown in the laboratory and from naturally-infected field-grown plants**

Authors: Pim Goossens, Kim Baremans, Marrit Alderkamp, Jordi C. Boshoven, Guido van den Ackerveken, Roeland L. Berendsen

**The following Supporting Information is available for this article:**

Supplementary Figures

Figure S1. **Bacterial community shifts in Viroflay phyllospheres upon *Pe* treatment at additional laboratories.**

Figure S2. ***Pe* abundances in naturally-infected field-grown Viroflay plants**

Figure S3. **Relative abundances of ASVs that were significantly enriched in *Pe*-infected field grown Viroflay leaves at Warmenhuizen.**

Figure S4. **Relative abundances of ASVs that were significantly enriched in *Pe*-infected field grown Viroflay leaves at Warmenhuizen.**

Supplementary Tables

Table S1. **PERMANOVA based on Bray-Curtis dissimilarities between all Viroflay and Caladonia leaf samples per treatment.**

Table S2. **Pairwise PERMANOVA analyses based on Bray-Curtis dissimilarities between samples, based on treatment (untreated, *Pe*10, *Pe*11, or *Pe*14) per cultivar per *Pe* inoculation method.**

Table S3. **PERMANOVA based on Bray-Curtis dissimilarities between all untreated and *Pe* water-inoculated Viroflay and Caladonia leaf samples.**

Table S4. **ASVs enriched in Pe-infected field grown Viroflay leaves at Warmenhuizen.**

Table S5. **ASVs enriched in Pe-infected field grown Viroflay leaves at Andijk.**

Table S6. **Primers used in this study.**

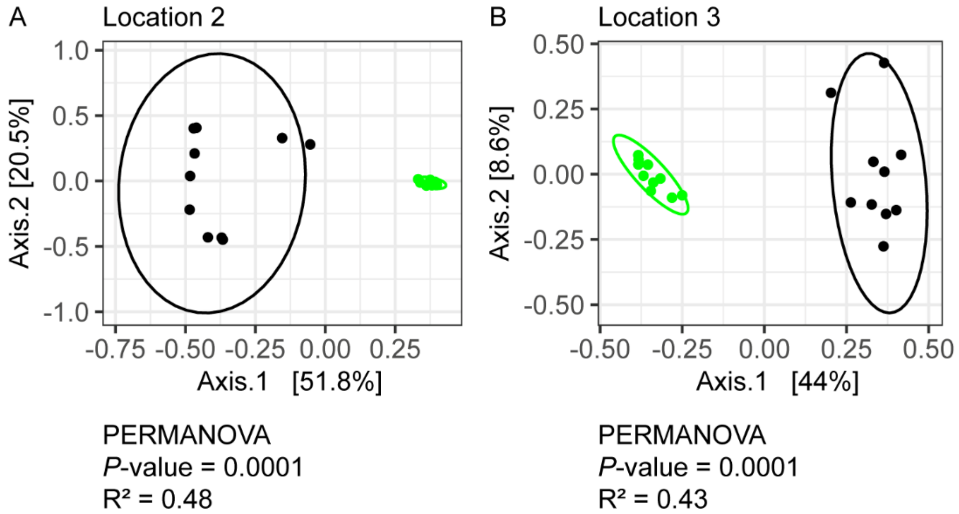


Figure S1. **Bacterial community shifts in Viroflay phyllospheres upon *Pe* treatment at additional laboratories.** (**A**, **B**) PCoA ordination plots based on Bray-Curtis dissimilarities of untreated and *Pe*14 water-inoculated Viroflay plants at (**A**) Location 2 and (**B**) Location 3. Symbol colors represent untreated (black symbols) and *Pe*14-2-inoculated (green symbols) plants at Location 2; and untreated (black symbols)and *Pe*14-3-inoculated (green symbols) plants at Location 3. Text below the graphs show PERMANOVA results, per location, for the comparison between *Pe*-inoculated plants and untreated plants based on 9999 permutations.

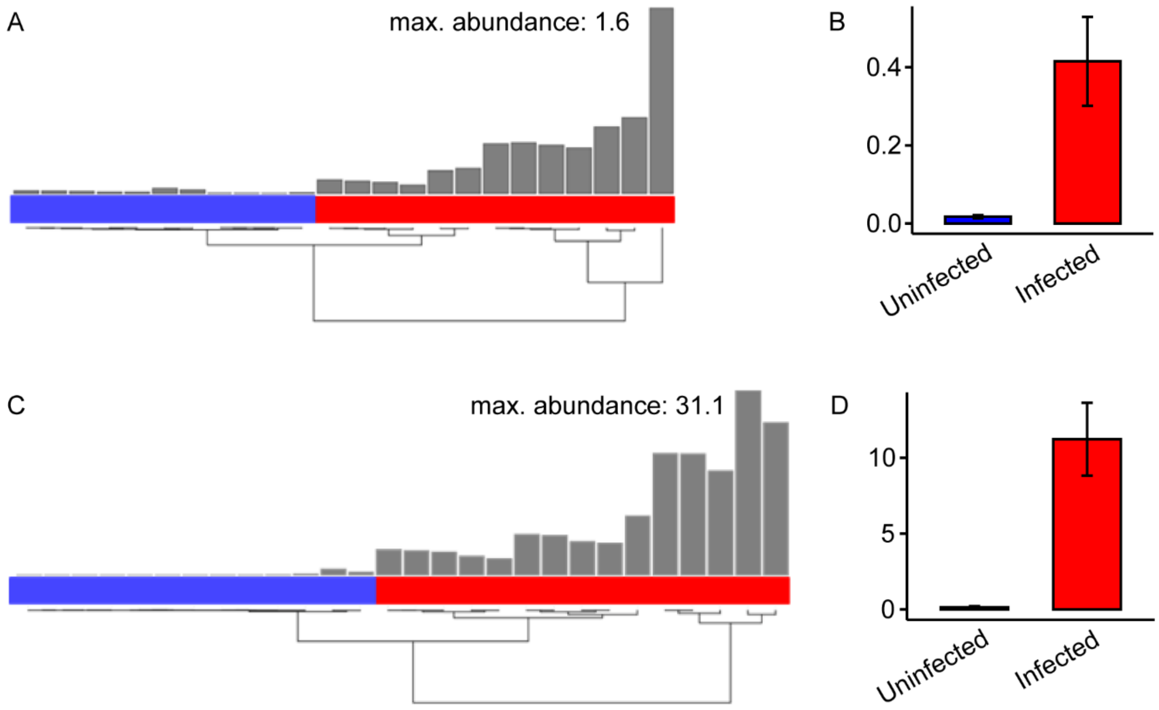


Figure S2. ***Pe* abundances in naturally-infected field-grown Viroflay plants** near (**A**, **B**) Warmenhuizen and (**C**, **D**) Andijk based on qPCR quantification of *Pe* genomic *ACTIN*, normalized by spinach genomic *ACTIN*. (**A**, **C**) Hierarchal clustering of samples based on *Pe* abundance and classification of samples into 'infected' (red labels) and 'uninfected' (blue labels). (**B**, **D**) Bar graphs of *Pe* mean abundance in samples from the respective fields (*N* as indicated in **A** and **C**). Error bars indicate standard error.

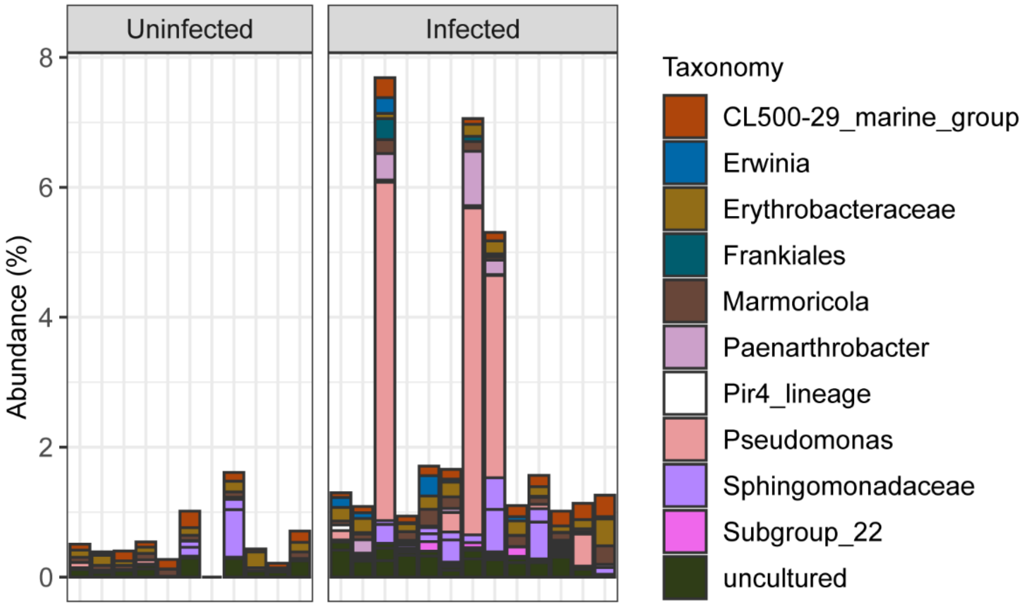


Figure S3. **Relative abundances of ASVs that were significantly enriched in *Pe*-infected field grown Viroflay leaves at Warmenhuizen.** Stacked bar graph showing the abundances of individual ASVs per leaf samples from the field at warmenhuizen. ASVs are colored based on Taxonomy as indicated on the right side of the graph.

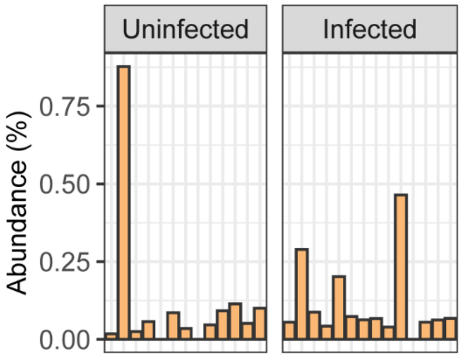


Figure S4. **Relative abundances of ASVs that were significantly enriched in *Pe*-infected field grown Viroflay leaves at Warmenhuizen.**

### Supplementary Tables

Table S1. **PERMANOVA based on Bray-Curtis dissimilarities between all Viroflay and Caladonia leaf samples per treatment.** $R^2$ is the proportion of variance attributed to the corresponding variable. F is the PERMANOVA pseudo F-statistic for the corresponding variable. Pr(>F) is the probability that the pseudo F-statistic of randomly permuted data (based on 9999 permutations) is larger than the pseudo F-statistic of the unpermuted data. *: *P*-value < 0.05.

| **Treatment** | **Tested variable** | **R2** | **F** | **Pr(>F)** |
|---|---|---|---|---|
| Untreated | Cultivar | 0.08077 | 1.5816 | 0.0247 * |
| Pe10_water | Cultivar | 0.10386 | 1.9702 | 0.0850 |
| Pe11_water | Cultivar | 0.08495 | 1.5782 | 0.1185 |
| Pe14_water | Cultivar | 0.05174 | 0.9822 | 0.3988 |
| Pe10_wind | Cultivar | 0.12014 | 2.4577 | 0.0002 *** |
| Pe11_wind | Cultivar | 0.07881 | 1.54 | 0.0746 |
| Pe14_wind | Cultivar | 0.06185 | 1.1867 | 0.2422 |

Table S2. **Pairwise PERMANOVA analyses based on Bray-Curtis dissimilarities between samples, based on treatment (untreated, *Pe*10, *Pe*11, or *Pe*14) per cultivar per *Pe* inoculation method.** Pairwise comparisons are indicated by 'Group 1' and 'Group 2'. $R^2$ is the proportion of variance attributed to the corresponding variable (i.e. treatment). Pr(>F) is the *fdr*-corrected probability that the pseudo F-statistic of randomly permuted data (based on 9999 permutations) is larger than the pseudo F-statistic of the unpermuted data for the sample set as indicated per row. *: *P*-value < 0.05; **: *P*-value < 0.01; ***: *P*-value < 0.001.

| **Group 1** | **Group 2** | **Cultivar** | **Pe_inoculation** | **$R^2$** | **Pr(>F)** |
|---|---|---|---|---|---|
| Untreated | Pe10 | Viroflay | Water | 0.32 | 0.0003 *** |
| Untreated | Pe11 | Viroflay | Water | 0.32 | 0.0001 *** |
| Untreated | Pe14 | Viroflay | Water | 0.37 | 0.0001 *** |
| Pe10 | Pe11 | Viroflay | Water | 0.46 | 0.0001 *** |
| Pe10 | Pe14 | Viroflay | Water | 0.44 | 0.0001 *** |
| Pe11 | Pe14 | Viroflay | Water | 0.21 | 0.0004 *** |
| Untreated | Pe10 | Caladonia | Water | 0.36 | 0.0002 *** |
| Untreated | Pe11 | Caladonia | Water | 0.29 | 0.0001 *** |
| Untreated | Pe14 | Caladonia | Water | 0.34 | 0.0001 *** |
| Pe10 | Pe11 | Caladonia | Water | 0.64 | 0.0001 *** |
| Pe10 | Pe14 | Caladonia | Water | 0.6 | 0.0001 *** |
| Pe11 | Pe14 | Caladonia | Water | 0.3 | 0.0001 *** |
| Untreated | Pe10 | Viroflay | Wind | 0.05 | 0.4112 |
| Untreated | Pe11 | Viroflay | Wind | 0.07 | 0.1563 |
| Untreated | Pe14 | Viroflay | Wind | 0.09 | 0.014 * |
| Pe10 | Pe11 | Viroflay | Wind | 0.07 | 0.116 |
| Pe10 | Pe14 | Viroflay | Wind | 0.09 | 0.0084 ** |
| Pe11 | Pe14 | Viroflay | Wind | 0.09 | 0.0202 * |
| Untreated | Pe10 | Caladonia | Wind | 0.07 | 0.093 |
| Untreated | Pe11 | Caladonia | Wind | 0.06 | 0.3317 |
| Untreated | Pe14 | Caladonia | Wind | 0.06 | 0.3011 |
| Pe10 | Pe11 | Caladonia | Wind | 0.05 | 0.4433 |
| Pe10 | Pe14 | Caladonia | Wind | 0.07 | 0.175 |
| Pe11 | Pe14 | Caladonia | Wind | 0.06 | 0.3383 |

Table S3. **PERMANOVA based on Bray-Curtis dissimilarities between all untreated and *Pe* water-inoculated Viroflay and Caladonia leaf samples** (PERMANOVA formula: *Pe* isolate * Cultivar). $R^2$ is the proportion of variance attributed to the corresponding variable. F is the PERMANOVA pseudo F-statistic for the corresponding variable. Pr(>F) is the probability that the pseudo F-statistic of randomly permuted data (based on 9999 permutations) is larger than the pseudo F-statistic of the unpermuted data. ***: *P*-value < 0.001.

| **Variable** | **$R^2$** | **F** | **Pr(>F)** |
|---|---|---|---|
| Pe_isolate | 0.43534 | 19.5694 | 0.0001 *** |
| Cultivar | 0.01385 | 1.8672 | 0.0613 |
| Pe_isolate:Cultivar | 0.03174 | 1.4267 | 0.0875 |
| Residual | 0.51907 | | |
| Total | 1 | | |

Table S4. **ASVs enriched in Pe-infected field grown Viroflay leaves at Warmenhuizen.**

| ASV | Prevalence in Pe lab cultures | Average abundance (%) in uninfected | Average abundance (%) in infected |
|---|---|---|---|
| *Paenarthrobacter* 1e165 | 4/5 | 0.00 | 0.14 |
| *Pseudomonas* 5e37b | 2/5 | 0.02 | 1.11 |
| *uncultured* 61385 | 0 | 0.13 | 0.24 |
| *Erythrobacteraceae* 3f3ff | 0 | 0.10 | 0.19 |
| CL500-29_marine_group 7ced5 | 0 | 0.11 | 0.17 |
| *Sphingomonadaceae* b83e5 | 0 | 0.08 | 0.15 |
| *Marmoricola* eb644 | 0 | 0.07 | 0.14 |
| *Sphingomonadaceae* e4f82 | 0 | 0.02 | 0.10 |
| *Erwinia* cd2d6 | 0 | 0.00 | 0.07 |
| uncultured a8a3f | 0 | 0.01 | 0.04 |
| *Frankiales* d0f2c | 0 | 0.00 | 0.04 |
| *Subgroup*_22 9ea58 | 0 | 0.00 | 0.03 |
| uncultured 4ef1d | 0 | 0.00 | 0.03 |
| uncultured 47ea5 | 0 | 0.01 | 0.02 |
| Pir4_lineage 2cf7c | 0 | 0.00 | 0.02 |
| uncultured 93752 | 0 | 0.00 | 0.02 |

Table S5. **ASVs enriched in Pe-infected field grown Viroflay leaves at Andijk.**

| ASV | Prevalence in Pe lab cultures | Average abundance (%) in uninfected | Average abundance (%) in infected |
|---|---|---|---|
| *Rhodococcus* 9c8fe | 5/5 | 0.12 | 0.11 |
| *Microbacteriaceae* 8fc0f | 2/5 | 2.08 | 7.85 |
| *Sphingomonas* f359d | 1/5 | 3.79 | 6.24 |
| *Pseudarthrobacter* d1316 | 0 | 5.51 | 30.62 |
| *Pseudarthrobacter* c1d8f | 0 | 5.16 | 8.60 |
| *Exiguobacterium* 41452 | 0 | 3.56 | 6.58 |
| *Massilia* c4958 | 0 | 0.05 | 0.70 |
| *Microbacteriaceae* 1ef23 | 0 | 0.07 | 0.53 |
| *Microbacteriaceae* 498ad | 0 | 0.16 | 0.37 |
| *Lysinimonas* 87d66 | 0 | 0.40 | 0.28 |
| *Bradyrhizobium* 9db28 | 0 | 0.29 | 0.16 |
| *Pseudarthrobacter* 1129c | 0 | 0.01 | 0.11 |
| *Pseudarthrobacter* 23c8a | 0 | 0.02 | 0.09 |
| *Clavibacter* 4d837 | 0 | 0.02 | 0.09 |
| *Paenarthrobacter* 844b8 | 0 | 0.01 | 0.08 |
| *Oxalobacteraceae* 2fccb | 0 | 0.04 | 0.06 |
| *Clavibacter* f9e85 | 0 | 0.01 | 0.06 |
| *Pseudarthrobacter* 6cfd7 | 0 | 0.01 | 0.05 |
| *Massilia* 11787 | 0 | 0.02 | 0.05 |
| *Rhodococcus* 43515 | 0 | 0.03 | 0.04 |
| *Paracoccus* 63086 | 0 | 0.03 | 0.04 |
| *Hymenobacter* 7b59c | 0 | 0.02 | 0.03 |
| *Paenarthrobacter* cb322 | 0 | 0.00 | 0.02 |
| *Frondihabitans* 7a772 | 0 | 0.01 | 0.02 |
| *uncultured* 3d168 | 0 | 0.00 | 0.02 |
| *uncultured* 00234 | 0 | 0.00 | 0.01 |
| *Blastococcus* a7ce1 | 0 | 0.00 | 0.01 |
| *Noviherbaspirillum* 793f8 | 0 | 0.00 | 0.01 |
| *Nocardioides* 70191 | 0 | 0.01 | 0.01 |
| uncultured 12d80 | 0 | 0.01 | 0.01 |
| uncultured 7d503 | 0 | 0.00 | 0.01 |

Table S6. **Primers used in this study.**

| **Name** | **Sequence (5' - 3')** | **Used for** | **Comment** |
|---|---|---|---|
| Actin_Pe_Fw | TCCAGGTGTTGGTGAACGTA | qPCR quantification of *Pe* | This study |
| Actin_Pe_Rv | TAGCGACGACAAGATGGAG | qPCR quantification of *Pe* | This study |
| Actin_Spinach_Fw | TCCGTGCAGGTATTGTGCTG | qPCR quantification of *Pe* | Duressa *et al.* (2012) |
| Actin_Spinach_Rv | AGCAAGGTCGAGACGAAGG | qPCR quantification of *Pe* | Duressa *et al.* (2012) |
| NGS1-16s-N701 | TCGTCGGCAGCGTCAGATGTGTATAAGAGACAGTCGCCTTACCTGTGGCCTACGGGNGGCWGCAG | PCR1 - Illumina 16S rDNA library preparation | Goossens *et al.* (2023) |
| NGS1-16s-N702 | TCGTCGGCAGCGTCAGATGTGTATAAGAGACAGCTAGTACGGAGTGGCCTACGGGNGGCWGCAG | PCR1 - Illumina 16S rDNA library preparation | Goossens *et al.* (2023) |
| NGS1-16s-N703 | TCGTCGGCAGCGTCAGATGTGTATAAGAGACAGTTCTGCCTTGCGACCTACGGGNGGCWGCAG | PCR1 - Illumina 16S rDNA library preparation | Goossens *et al.* (2023) |
| NGS1-16s-N704 | TCGTCGGCAGCGTCAGATGTGTATAAGAGACAGGCTCAGGAATGACCTACGGGNGGCWGCAG | PCR1 - Illumina 16S rDNA library preparation | Goossens *et al.* (2023) |
| NGS1-16s-N705 | TCGTCGGCAGCGTCAGATGTGTATAAGAGACAGAGGAGTCCCGACCTACGGGNGGCWGCAG | PCR1 - Illumina 16S rDNA library preparation | Goossens *et al.* (2023) |
| NGS1-16s-N706 | TCGTCGGCAGCGTCAGATGTGTATAAGAGACAGCATGCCTACGACCTACGGGNGGCWGCAG | PCR1 - Illumina 16S rDNA library preparation | Goossens *et al.* (2023) |
| NGS1-16s-N707 | TCGTCGGCAGCGTCAGATGTGTATAAGAGACAGGTAGAGAGGTCCTACGGGNGGCWGCAG | PCR1 - Illumina 16S rDNA library preparation | Goossens *et al.* (2023) |
| NGS1-16s-N708 | TCGTCGGCAGCGTCAGATGTGTATAAGAGACAGCCTCTCTGGTCCTACGGGNGGCWGCAG | PCR1 - Illumina 16S rDNA library preparation | Goossens *et al.* (2023) |
| NGS1-16s-N709 | TCGTCGGCAGCGTCAGATGTGTATAAGAGACAGAGCGTAGCTCCTACGGGNGGCWGCAG | PCR1 - Illumina 16S rDNA library preparation | Goossens *et al.* (2023) |
| NGS1-16s-N710 | TCGTCGGCAGCGTCAGATGTGTATAAGAGACAGCAGCCTCGTCCTACGGGNGGCWGCAG | PCR1 - Illumina 16S rDNA library preparation | Goossens *et al.* (2023) |
| NGS1-16s-N711 | TCGTCGGCAGCGTCAGATGTGTATAAGAGACAGTGCCTCTTCCTACGGGNGGCWGCAG | PCR1 - Illumina 16S rDNA library preparation | Goossens *et al.* (2023) |
| NGS1-16s-N712 | TCGTCGGCAGCGTCAGATGTGTATAAGAGACAGTCCTCTACCCTACGGGNGGCWGCAG | PCR1 - Illumina 16S rDNA library preparation | Goossens *et al.* (2023) |
| NGS1-16s-N501 | GTCTCGTGGGCTCGGAGATGTGTATAAGAGACAGTAGATCGCCACTTCTGACTACHVGGGTATCTAATCC | PCR1 - Illumina 16S rDNA library preparation | Goossens *et al.* (2023) |
| NGS1-16s-N502 | GTCTCGTGGGCTCGGAGATGTGTATAAGAGACAGCTCTCTATTTCTCTGACTACHVGGGTATCTAATCC | PCR1 - Illumina 16S rDNA library preparation | Goossens *et al.* (2023) |
| NGS1-16s-N503 | GTCTCGTGGGCTCGGAGATGTGTATAAGAGACAGTATCCTCTACTCAGACTACHVGGGTATCTAATCC | PCR1 - Illumina 16S rDNA library preparation | Goossens *et al.* (2023) |
| NGS1-16s-N504 | GTCTCGTGGGCTCGGAGATGTGTATAAGAGACAGAGAGTAGAGATAGACTACHVGGGTATCTAATCC | PCR1 - Illumina 16S rDNA library preparation | Goossens *et al.* (2023) |
| NGS1-16s-N505 | GTCTCGTGGGCTCGGAGATGTGTATAAGAGACAGGTAAGGAGCTAGACTACHVGGGTATCTAATCC | PCR1 - Illumina 16S rDNA library preparation | Goossens *et al.* (2023) |
| NGS1-16s-N506 | GTCTCGTGGGCTCGGAGATGTGTATAAGAGACAGACTGCATATCGACTACHVGGGTATCTAATCC | PCR1 - Illumina 16S rDNA library preparation | Goossens *et al.* (2023) |
| NGS1-16s-N507 | GTCTCGTGGGCTCGGAGATGTGTATAAGAGACAGAAGGAGTAAGACTACHVGGGTATCTAATCC | PCR1 - Illumina 16S rDNA library preparation | Goossens *et al.* (2023) |
| NGS1-16s-N508 | GTCTCGTGGGCTCGGAGATGTGTATAAGAGACAGCTAAGCCTGACTACHVGGGTATCTAATCC | PCR1 - Illumina 16S rDNA library preparation | Goossens *et al.* (2023) |
| S501 | AATGATACGGCGACCACCGAGATCTACACTAGATCGCTCGTCGGCAGCGTC | PCR2 - Illumina 16S rDNA library preparation | Illumina |
| S502 | AATGATACGGCGACCACCGAGATCTACACCTCTCTATTCGTCGGCAGCGTC | PCR2 - Illumina 16S rDNA library preparation | Illumina |
| S503 | AATGATACGGCGACCACCGAGATCTACACTATCCTCTTCGTCGGCAGCGTC | PCR2 - Illumina 16S rDNA library preparation | Illumina |
| S504 | AATGATACGGCGACCACCGAGATCTACACAGAGTAGATCGTCGGCAGCGTC | PCR2 - Illumina 16S rDNA library preparation | Illumina |
| S505 | AATGATACGGCGACCACCGAGATCTACACGTAAGGAGTCGTCGGCAGCGTC | PCR2 - Illumina 16S rDNA library preparation | Illumina |
| S506 | AATGATACGGCGACCACCGAGATCTACACACTGCATATCGTCGGCAGCGTC | PCR2 - Illumina 16S rDNA library preparation | Illumina |
| S507 | AATGATACGGCGACCACCGAGATCTACACAAGGAGTATCGTCGGCAGCGTC | PCR2 - Illumina 16S rDNA library preparation | Illumina |
| S508 | AATGATACGGCGACCACCGAGATCTACACCTAAGCCTTCGTCGGCAGCGTC | PCR2 - Illumina 16S rDNA library preparation | Illumina |
| N701 | CAAGCAGAAGACGGCATACGAGATTCGCCTTAGTCTCGTGGGCTCGG | PCR2 - Illumina 16S rDNA library preparation | Illumina |
| N702 | CAAGCAGAAGACGGCATACGAGATCTAGTACGGTCTCGTGGGCTCGG | PCR2 - Illumina 16S rDNA library preparation | Illumina |
| N703 | CAAGCAGAAGACGGCATACGAGATTTCTGCCTGTCTCGTGGGCTCGG | PCR2 - Illumina 16S rDNA library preparation | Illumina |
| N704 | CAAGCAGAAGACGGCATACGAGATGCTCAGGAGTCTCGTGGGCTCGG | PCR2 - Illumina 16S rDNA library preparation | Illumina |
| N705 | CAAGCAGAAGACGGCATACGAGATAGGAGTCCGTCTCGTGGGCTCGG | PCR2 - Illumina 16S rDNA library preparation | Illumina |
| N706 | CAAGCAGAAGACGGCATACGAGATCATGCCTAGTCTCGTGGGCTCGG | PCR2 - Illumina 16S rDNA library preparation | Illumina |
| N707 | CAAGCAGAAGACGGCATACGAGATGTAGAGAGGTCTCGTGGGCTCGG | PCR2 - Illumina 16S rDNA library preparation | Illumina |
| N708 | CAAGCAGAAGACGGCATACGAGATCCTCTCTGGTCTCGTGGGCTCGG | PCR2 - Illumina 16S rDNA library preparation | Illumina |
| N709 | CAAGCAGAAGACGGCATACGAGATAGCGTAGCGTCTCGTGGGCTCGG | PCR2 - Illumina 16S rDNA library preparation | Illumina |
| N710 | CAAGCAGAAGACGGCATACGAGATCAGCCTCGGTCTCGTGGGCTCGG | PCR2 - Illumina 16S rDNA library preparation | Illumina |
| N711 | CAAGCAGAAGACGGCATACGAGATTGCCTCTTGTCTCGTGGGCTCGG | PCR2 - Illumina 16S rDNA library preparation | Illumina |
| N712 | CAAGCAGAAGACGGCATACGAGATTCCTCTACGTCTCGTGGGCTCGG | PCR2 - Illumina 16S rDNA library preparation | Illumina |
| pPNA | GGCTCAACCCTGGACAG | PCR1 PCR clamp | Lundberg *et al.* (2013) |
| mPNA | GGCAAGTGTTCTTCGGA | PCR1 PCR clamp | Lundberg *et al.* (2013) |
